# Breaking barriers in the sensitive and accurate mass determination of large DNA plasmids by mass photometry

**DOI:** 10.1101/2025.03.25.645154

**Authors:** Eduard H.T.M. Ebberink, Evolène Deslignière, Alisa Ruisinger, Markus Nuebel, Marco Thomann, Albert J.R. Heck

## Abstract

DNA plasmids (pDNA) are essential for gene cloning and protein expression, whereby engineered plasmids serve as vectors to insert foreign DNA into host cells, enabling mass production of proteins and vaccines. Furthermore, pDNA is used in CRISPR-based gene editing, RNA therapeutics, and DNA vaccines. Due to the rapidly increasing use and application of a wide variety of pDNA, analytical methods to characterize their key attributes are vital. Because of their high molecular weight, accurate and fast mass analyses of pDNA, as a measure of quality control, is rather challenging. Here we explore mass photometry (MP) to analyze pDNAs and find that it completely fails using standard procedures as developed for MP on proteins, with masses underestimated by 30-40%. Even though the landing of pDNA during MP analysis can be improved by using coated glass slides, the large dsDNA particles diffract light beyond the diffraction limit, rendering most landing events unusable. To overcome these issues, we introduce a fast (30 s) and simple protocol to convert dsDNA particles rapidly into ssDNA-like particles just prior to analysis and show that these particles behave nearly perfect for MP. Using this protocol accurate and correct masses of pDNAs can be obtained by MP, with values within 1-3% of the expected mass. Using this protocol, MP can be used to mass analyze pDNA constructs from 1 to 15 MDa, suggesting that this approach may be widely adopted within academia and biopharma for essentially all plasmids.

## Introduction

Purified DNA plasmids are nowadays indispensable in preparing gene therapy and vaccine products.^1,2^ Whether in generating the genetic cargo, the protein-based capsid/carrier system, or in producing a completely protein-based pharmaceutical, an intact DNA plasmid (pDNA) usually forms the basis of gene transcription and, subsequently, protein expression. Because defined, purified pDNA acts as the template for protein expression, these plasmids are essential for use in biomolecular and pharmaceutical research, development and clinical applications. In all such studies and applications it is critical that the integrity and quality of pDNA can be accurately assessed on a regular and time- and cost-efficient basis.^3^

Besides a pivotal role in pharmaceutical manufacturing, a recent exciting development in the direct use of pDNA is as a vaccine agent, using delivery of ‘naked’ DNA to the human body. Vaccination with pDNA has gained traction with the approval of the first pDNA-based vaccine against SARS-CoV-2.^4^ Since the early 1990s, the idea of vaccination by pDNA-driven protein expression, which elicits a broad (B-cell and T-cell) immune response, has been explored.^5-8^ And, up to date, DNA vaccines remain a promising avenue for immunization, including for cancer therapy.^9,10^ DNA-based vaccines hold several intrinsic advantages, as (i) pDNA is highly stable (also at room temperature), (ii) relatively cost-efficient to produce, and (iii) easy to mutationally modify (e.g., to respond to seasonal influenza). Notwithstanding this great potential, pDNA has thus far only had limited clinical applications, mainly due to the significant challenge of efficiently delivering naked DNA to antigen-presenting cells and the cell nucleus in general.^11^ Despite ongoing developments in cell-free plasmid manufacturing (e.g., Doggybone^TM^ DNA) and focus on more efficient DNA delivery methods,^12-14^ the amount of clinical-grade pDNA needed as dosage is anticipated to be high.^2,10,15^ Therefore, analytic tools to better characterize and quality-control large intact DNA constructs need to be co-developed.^3^

Conventionally, a plasmid’s integrity is determined by size exclusion-based techniques such as gel filtration, agarose gel electrophoresis, or capillary electrophoresis.^16^ To further characterize clinical-grade pDNAs, the genetic sequence is generally assessed by methods using Sanger-, next-generation-, or nanopore-sequencing. While being the standard for many years, electrophoresis and sequencing runs can be costly, time-consuming or laborious and are not ideal for high-throughput applications, often desirable in biopharmaceutical research and development.

In recent years, mass photometry (MP) has become an effective method for fast and sensitive mass determination of biomolecules, although most applications have been targeting proteins and/or protein complexes. In MP, using interferometric microscopy, the mass of a single (protein) particle can be determined by the light it scatters once it lands on a glass surface and interferes with the reflected laser light.^17,18^ The scattering-induced interference with accompanied contrast value is proportionate to the mass of the particle. This single-particle analysis is fast, requires almost no sample preparation (e.g., label-free) and only a limited amount of material. As it is becoming an essential tool within biological research, MP has been applied to mostly protein-based samples. The application of MP for DNA constructs has been modestly explored, only in a few instances has pure DNA been assessed by MP.^19-23^ In these studies, MP was shown to be a viable method for detecting small DNA fragments.^19^ With additional adjustments from protein-based measurements (surface modification with 3-aminopropyltriethoxysilane (APTES) and applying an appropriate DNA calibration), the mass and length could be determined for DNA constructs up to 2000 base pairs (bps). However, already with these relatively small DNA fragments (compared to plasmids), the size of these DNA particles gets close to the diffraction limit, which affects contrast determination and, thus accurate mass analysis.^19,23^ Previously, also we observed odd-shaped landing events of the pBR322 DNA plasmid on an APTES-coated glass surface.^24^ With pBR322’s persistence length extending beyond the diffraction limit, the point source signal cannot be described anymore as a 2D (Gaussian function-derived) circular blur/dot by the interferometric point-spread function (PSF).^23,25^ As a result, with landing events becoming less circular, practically all signals become useless for mass analysis during conventional MP data processing.

This study presents a fast and effective way to deal with the plasmid length and the MP diffraction limit. By exploiting distinct DNA morphologies, especially considering double-stranded *versus* single-stranded conformations, we present a fast protocol based on acid-induced conversion and use it to successfully measure accurate masses (i.e., within ∼3%) of plasmids up to ∼7250 base pairs, even on unmodified glass surfaces. By introducing this cheap and efficient conformational conversion protocol, MP can be used to accurately measure practically each DNA plasmid, as demonstrated here for constructs from 1 to 15 MDa.

## Results

### Plasmid samples studied

To evaluate the challenges in, and potential of, MP to characterize pDNA we targeted four different large-sized dsDNA plasmids, namely pUC18 (2686 bps), pBR322 (4361 bps), ΦX174 (5386 bps) and M13mp18 (7249 bps). Details on these samples are summarized in Table 1. In previous work from others and our group, it was already demonstrated that DNA molecules often behave very differently in MP when compared to their (molecular weight-alike) protein counterparts.^19,23,24^ First, landing on non-modified glass slides, normally used in MP, is not ideal for negatively charged DNA molecules. Indeed, as expected, no signal was observed when using conventional glass slides to measure the abovementioned pDNA constructs. Derivatizing the glass slides with chemicals that effectively “charge” the surface (for instance, polyLys or APTES), is needed to alleviate these issues.^19,24^ Therefore, also here we started first by using APTES-coated glass slides to perform MP on the dsDNA plasmids (Table 1, using standard MP measurement conditions). On APTES-coated slides, we could detect several pDNA landing events. However, when applying the standard protein-based calibration curve the processed mass values do not at all match the theoretical values, with deviations ranging from 30-40 % (see Table 1). Generally, mass calibration in MP is done by using a series of protein assemblies of known molecular weight. However, the molecular polarizability of globular proteins *versus* linear nucleotide-based particles can differ significantly.^20^ Therefore, such a calibration curve fails when analyzing RNA or DNA molecules. Measuring pDNA thus requires a different approach.

**Table 1:** Plasmids analyzed in this study and masses measured by MP. Provided in the first three columns are the names of the commercially available plasmids, their nucleotide sequence lengths and theoretical masses. In the next four columns are depicted the measured masses of these constructs by MP, using different conditions and protocols.

| | Nucleotides | Theo. mass (MDa)<sup>a</sup> | Standard MP (mass ± FWHM in MDa) | | DNA optimized MP (mass ± FWHM in MDa) | |
| --- | --- | --- | --- | --- | --- | --- |
|  |  |  | APTES coated | Glass<sup>b</sup> | APTES coated | Glass |
| pUC18 | 5372 | 1.66 | 1.19 ± 0.29 | ND | 1.74 ± 0.13 | 1.66 ± 0.14 |
| pBR322 | 8722 | 2.69 | 1.79 ± 0.41 | ND | 2.81 ± 0.17 | 2.72 ± 0.18 |
| ΦX174 | 10772 | 3.33 | 2.14 ± 0.49 | ND | 3.42 ± 0.17 | 3.36 ± 0.21 |
| M13mp18 | 14498 | 4.47 | 2.61 ± 0.45 | ND | 4.66 ± 0.30 | 4.61 ± 0.22 |
| p8064 | 8064 | 2.49 | 2.32 ± 0.17 | 2.51 ± 0.21 | 2.45 ± 0.17 | 2.52 ± 0.14 |

<sup>a</sup>Theoretical mass calculated from the DNA sequence. <sup>b</sup>ND, not detected.

### Analysis of double-stranded pDNA mass photometry is constrained by the diffraction limit

Critically viewing the landing events in MP, we noticed that the particles landing on an APTES-coated surface resulted in both circular and oval-shaped features, in line with what we reported previously (Fig. 1A).^24^ In MP, these non-circular shaped signals, do not fall within the expected point-spread function (PSF) of ideal landing events and are usually discarded when standard filters are applied in the analysis of the landing events. Using the standard analysis protocol a contrast histogram with a Gaussian distribution is obtained with an average contrast value of ∼-0.0183 (Fig. 1B). Conversely, when all landing particles are considered regardless of their shape features, the average contrast signal appears to be underestimated (∼-0.0164, Fig. 1B). These latter non-ideal landing events do not fit the PSF and add lower-than-expected contrast signal to the distribution, as reflected by the low contrast shoulder observed in the contrast histogram (Fig. 1B). Taking such contrast difference into account, results in a substantial average mass shift of ∼185 kDa. The misalignment with the protein standard used for mass calibration was expected as lengthy and anisotropic dsDNA constructs have different polarizability compared to globular proteins.^19^ However, as the mass assignment is directly related to the contrast values of the particles, the underestimation of the contrast even with strict filtering in place leads to artificially, substantially lower mass values for all studied plasmids, which become apparent when constructing a high mass standard curve of the measured ds pDNA (Fig. 1C). Especially in the case of the larger ds pDNA, notably ΦX174 (5386 bps) and M13mp18 (7249 bps) this effect of the PSF misfitting is unmistakable. Because larger-sized plasmids suffer from a more substantial misfitting effect, this tends to force the curve to become non-linear and, thus, not useful as a standard.

**Figure 1:**
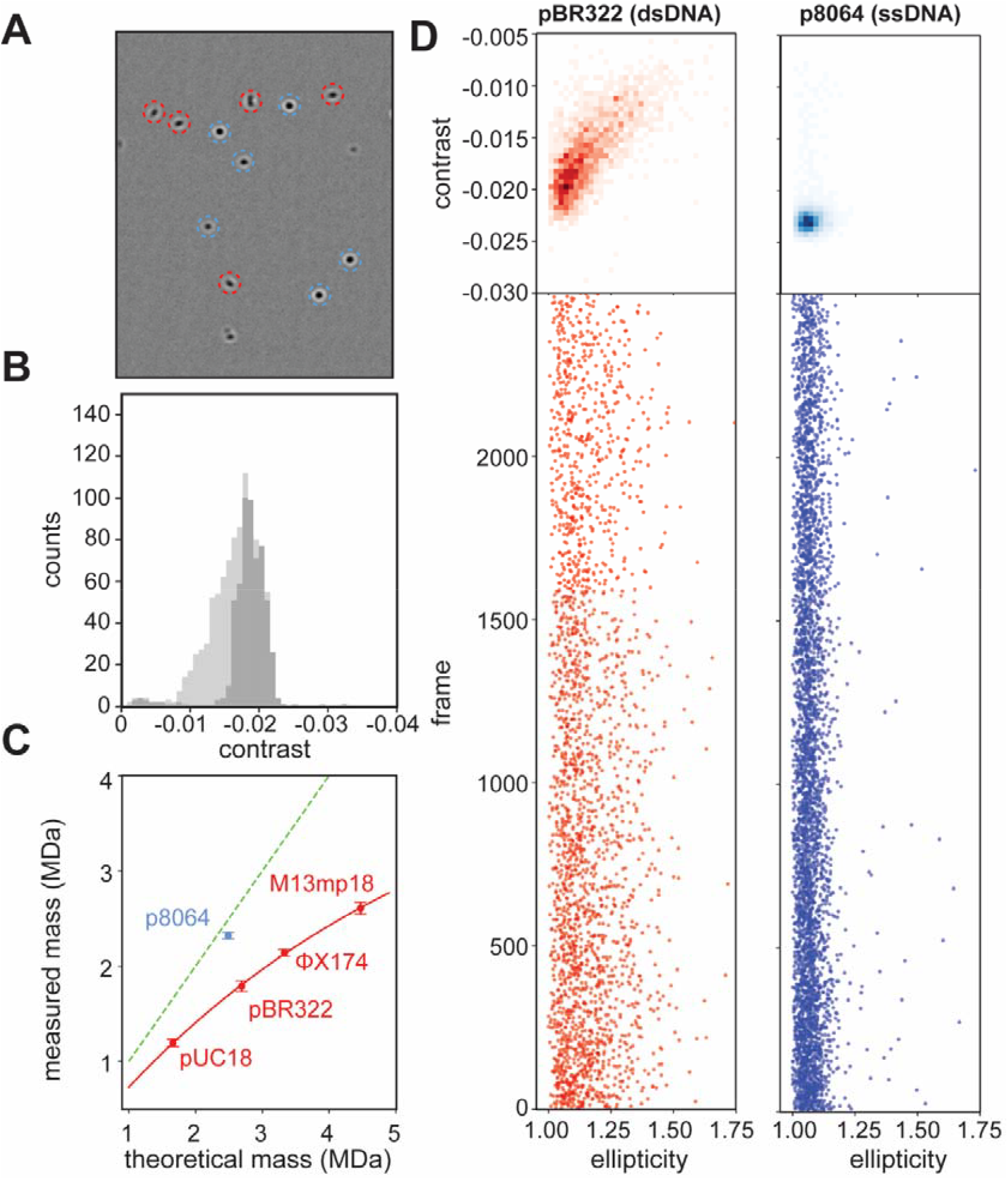
Double stranded plasmids display non-ideal behavior in mass photometry. **A)** Video frame of a typical pBR322 recording. Next to the desired circular (blue) landing events also many non-circular, elliptic (red) contrast signals are observed. **B)** Contrast histogram of landed pBR322 particles. When non-circular landing events are discarded based on their poor point-spread function (PSF) fit, the resulting contrast histogram is Gaussian (dark grey). Processing of all particles reveals a broader, lower-skewed distribution (light grey). **C)** Plot of expected *versus* extracted masses of pDNAs. The green dashed line represents the ideal match, in red are the data for the dsDNA plasmids and in light blue the data for the ssDNA. The red line through the data for the dsDNA curves off at higher masses, but also clearly shows that the measured masses of all dsDNA plasmids are far below the expected masses, while the mass of the ssDNA matches the expected mass. The depicted error bars represent the standard deviations (n>3). **D)** Contrast and ellipticity histograms of dsDNA pBR322 and ssDNA p8064, revealing the substantially lower number of non-circular landing events for the ssDNA

In contrast, when applying MP under identical conditions on a ssDNA plasmid p8064, of similar size as pBR322 (8064 nt), the contrast-to-mass value matches much better that of protein assemblies with the same mass (Fig. 1C, Table 1). For the ssDNA plasmid the number of non-ideal, oval-shaped signals in MP is substantially lower in the recorded images. The landing events now contain mostly circular shapes, fitting well the PSF (Fig. 1C and 1D). That the refractive index of structurally less ordered ssDNA matches more closely that of proteins was already known,^19^ but also, the shape of the p8064 MP signals appears more circular than dsDNA and appears to nicely fit the PSF. In addition, the ssDNA particles appear to interact well with the non-modified glass slides and can thus be mass analyzed without coating the glass with APTES (see Table 1). From this analysis, we extracted the hypothesis that unfolding and breaking dsDNA plasmids into ssDNA-like particles might be beneficial for the accurate mass analysis of the pDNA by MP.

### Transformation of double-stranded DNA particles into single-stranded-like particles enables accurate mass measurement by mass photometry

As the ssDNA plasmid particles behaved extremely well in MP, we sought means to convert dsDNA particles into ssDNA-like particles just prior to MP analysis. To test whether we could convert the dsDNA plasmid of pBR322 into ssDNA-like particles, we incubated the dsDNA with various amounts of formic acid (FA). We expected that the low pH (< 2) would denature the dsDNA, possibly resulting in ssDNA-like assemblies, ideally without falling apart into fragments. Following incubation with FA, we first attempted to separate the ssDNA-like particles from residual dsDNA plasmids using C18 reversed-phase high-pressure liquid chromatography (RP-HPLC), as it has been described that ssDNA is expected to be retained longer on such column material (Fig. S1A).^26^ With increased FA incubation time, we did observe an increasing population of particles eluting as a second peak, much later than the original dsDNA particles (Fig.2A). Incubation with a higher percentage of FA also enhanced the formation of this late-eluting population of pBR322 particles. The pUC18 plasmid behaved similarly upon the addition of FA (Fig. S1B). When sampling by MP the early eluting fraction (∼ 25 minutes, peak I) of FA-treated pBR322, we observed in the video frames oval-shaped landing events, as seen before for the dsDNA plasmids. These particles have an apparent, average (experimental) mass of ∼1.9 MDa, resembling the measurement of double-stranded pBR322 directly studied by MP (Fig. 1, 2B and 2C). For the late eluting fraction (∼ 29 minutes, peak II) no oval-shaped landing events were observed in MP and the extracted average mass of the particles (∼2.6 MDa) was much closer to the theoretical mass of pBR322 (2.69 MDa). Because particles from the late elution peak, generated by FA treatment, displayed similar characteristics and retention time as the previously measured ssDNA p8064, we concluded that we successfully prepared a ssDNA-like conformer out of the dsDNA pBR322 plasmids that in mass is still representative of the original dsDNA plasmid.

**Figure 2:**
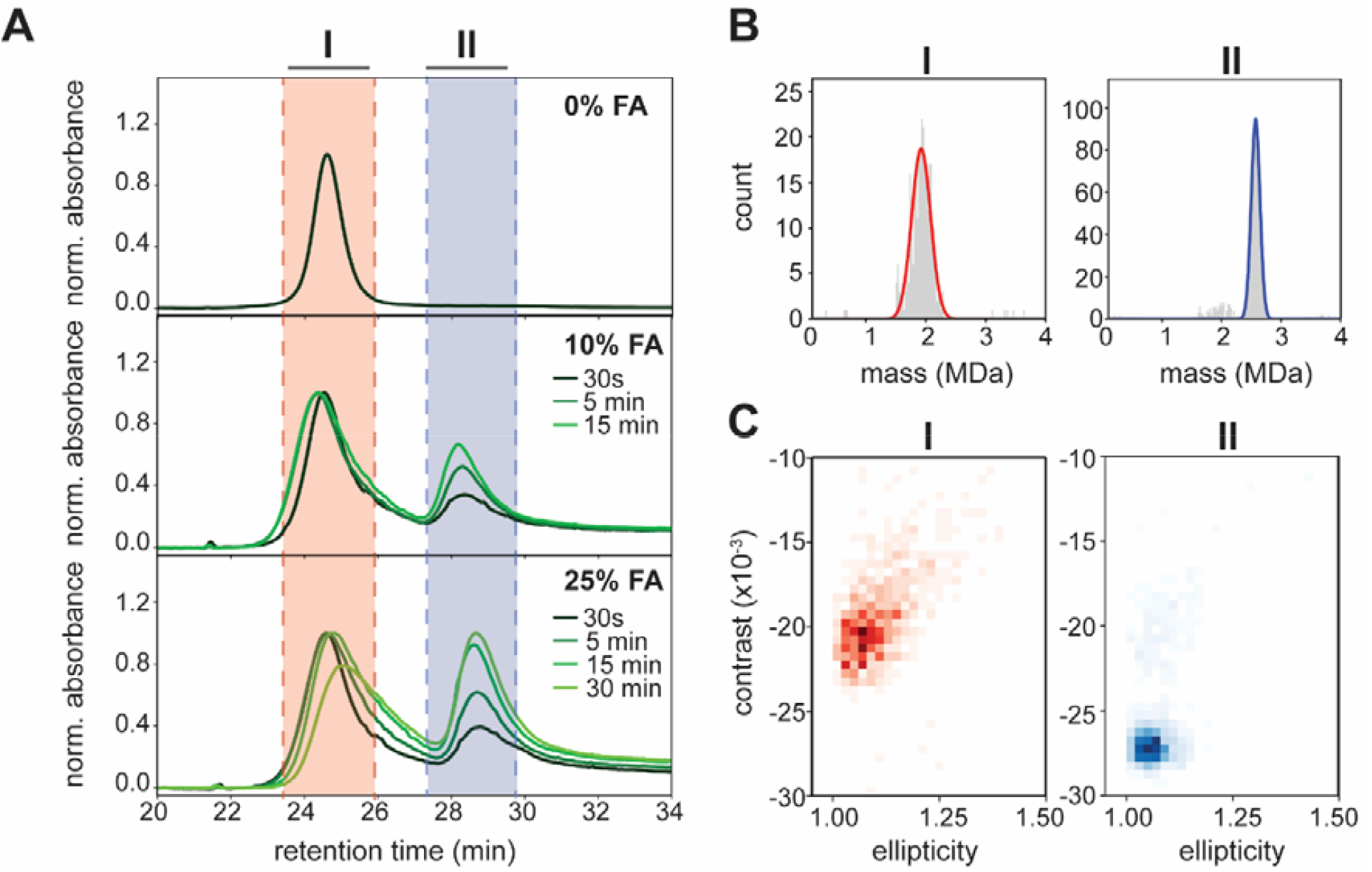
Incubating dsDNA plasmid particles in formic acid transforms them to ssDNA-like particles amendable for accurate and correct mass photometry measurement. **A)** Transformation of pBR322 plasmid to ssDNA-like particles by formic acid (FA) monitored by RP-HPLC. Displayed are overlays of the chromatograms of pBR322 incubated with different percentages of FA (i.e., 10 and 25%) for different time periods (from dark to light green: 30 s, 5 min, 15 min and 30 min). Untreated pBR322 elutes at ∼25 minutes (peak I, red shaded). FA incubation produces a new peak at ∼29 minutes (peak II, blue shaded), resembling single-stranded p8064 (see Fig. S1). Longer incubation and/or higher FA concentrations increase the amounts of ssDNA-like particles. **B)** Following HPLC separation, fractions taken from the elution peaks at 25 minutes and 29 minutes were analyzed by MP. Particles eluted at the 25-minute mark (peak I) led to similar data as for untreated pBR322 with many non-circular landing events, leading to incorrect lower masses (∼1.9 MDa). Particles taken from the 29-minute elution peak (peak II) displayed mostly circular shapes in MP. Moreover, the masses based on the extracted contrast values (∼2.6 MDa) now match closely the expected mass (2.69 MDa) of a pBR322 plasmid (see also Table 1) even when using a protein standard for MP mass calibration. **C)** The signals’ ellipticity was calculated for the fractions recorded in panel B. The 2D-histogram shows that native pBR322 from peak I contains many non-circular landing events while ssDNA-like pBR322 from peak II generates exclusively circular landing events.

### An efficient protocol for dsDNA to ssDNA-like particle conversion exploitable for mass photometry

To optimize the MP protocol for pDNA analysis we further explored the rapid conversion of dsDNA into ssDNA-like particles by using FA. Therefore, we next tested a short denaturation time of just 30 seconds with an MP-compatible concentration of pBR322 (12.5 ng/µL) and variable percentages of FA using the APTES-coated glass slides. Gradually increasing the amount of FA revealed the pathway of dsDNA to ssDNA-alike particle conversion (Fig. 3A). At a low FA concentration of 0.1%, a broad distribution of dsDNA particles is observed with an apparent average mass slightly shifted compared to the measured mass in the absence of FA (Table 1, Fig. 1C and 3A). When the FA concentration is increased, this most abundant peak gradually shifts towards higher masses, reaching a final sharp mass distribution with an average mass of ∼2.8 MDa at 10% FA. At 2.5% FA, part of the initial pBR322 dsDNA population splits into a single ssDNA segment (1x ssDNA) or dimers of such ssDNA segments (2x ssDNA), yielding particles with a mass corresponding to half and full pBR322 plasmids, respectively. Both these new populations flank the main dsDNA distribution, which is still present in this sample. The single ssDNA segment is only detected in our data when using 2.5% FA. Notably, when all the dsDNA pBR322 plasmids are converted into ssDNA-like assemblies (at 10% FA), a further increase of the FA concentration leads to the formation of higher mass multimers (Fig. 3B). For pBR322 we observe even pentamers and hexamers, with extracted masses of 13.7 and 16.2 MDa, although such large particles are typically expected to be beyond the higher mass limit of the used Samux^MP^ mass photometer. The extracted multimer masses are still quite close to the expected masses for such MDa assemblies. This DNA oligomerization, induced by high concentrations of FA, could also be reproduced for ssDNA p8064 (Fig. 3B). In a highly similar pattern, we detected p8064 oligomers up to pentamers and extracted accurate expected masses of ∼12.2 MDa. Therefore, we propose that such induced oligomer ladders of ssDNA aggregates may be used as (cheaper and more readily available) calibrants in MP. So far, in these measurements, we still used the APTES-coated glass surfaces, because we anticipated that these would improve the landing and binding of dsDNA molecules. At the same time, we observed that ssDNA p8064 worked equally well on non-modified glass slides. Therefore, now that we developed a fast protocol to convert dsDNA into ssDNA-like particles, we next explored whether we could simply use non-modified glass slides also for analysing these particles. Once incubated with FA, the converted dsDNA pBR322 particles indeed start to interact much better with the non-modified glass surface. Even at 0.1% FA, APTES coating seems no longer required to detect signals. However, at this lower FA concentration, the resulting landing events are relatively faint, yielding low contrast values, and thus, incorrect lower apparent masses (Fig. 3A). However, with a short FA incubation above 2%, the acquired data on the non-modified glass plates become alike those on the APTES coated slides (Fig. 3A). And therefore, we suggest that MP can be used to assess accurate masses of DNA plasmids, even when using non-modified glass surfaces.

**Figure 3:**
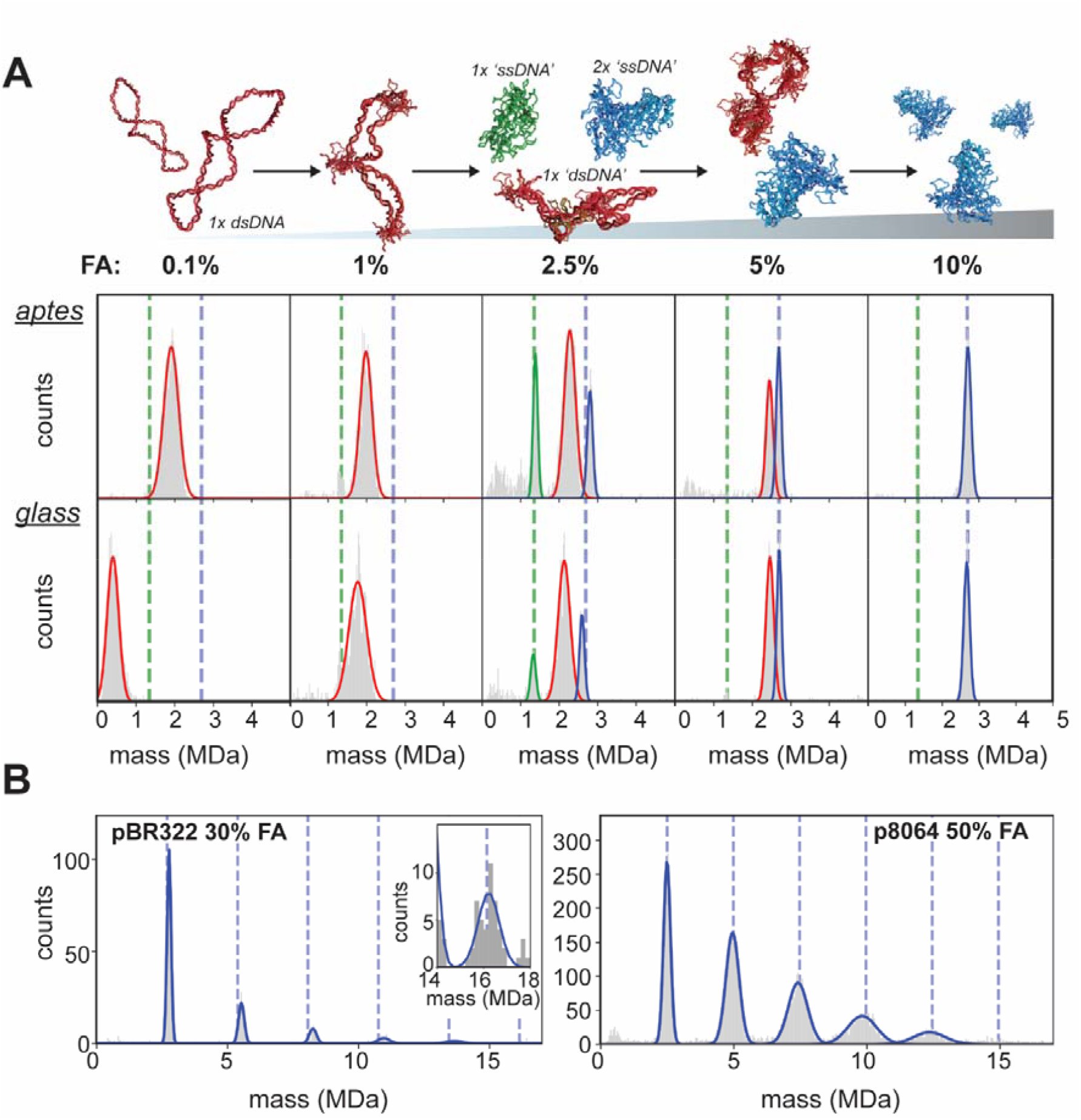
Monitoring formic acid induced transformation of dsDNA into ssDNA-like particles by mass photometry. **A)** All samples were incubated with FA for 30 s. In the top row, the FA-induced changes in dsDNA pBR322 at different concentrations of FA are analyzed by MP using APTES-coated slides, whereas, in the bottom row, non-modified glass slides were used. Due to an increase in contrast, the main population of pBR322 particles starts to shift towards a higher apparent mass, whereby the particles fragment into 1x ssDNA-like or 2x ssDNA-like pBR322 particles (mass indicated by respectively green and blue vertical, dashed lines). Using 10% FA, a very sharp distribution of particles is observed with an average accurate mass close to the theoretical mass off pBR322. In the mass histograms, the Gaussian fits of different populations are color-coded. **B)** At even higher FA concentrations (30% and 50%) the 2x ssDNA-like pBR322 particles start to multimerize, forming even up to hexamers with a molecular weight of ∼16 MDa. Multimer masses of pBR322 are indicted by blue dashed lines. The inset on the left shows the accumulated signal of several measurements. A similar oligomerization process is observed for ssDNA p8064 particles (right).

### DsDNA denaturation by FA improves mass accuracy and extends the range of detection

Optimizing the parameters with dsDNA pBR322, we found that a short (30 s) incubation step with 10% FA is optimal to obtain a single mass distribution in MP from which a mass can be extracted that corresponds well to the theoretical mass of the plasmid (Fig 3A). Next, we analyzed several other dsDNA plasmids and observed similar effects, namely (i) mostly circular landing events after incubation with FA and (ii) a shift towards higher contrast, resulting in accurate masses in MP (Fig. 4A and S2). After FA transformation of the dsDNA into ssDNA-like particles the inferred masses align very well with the theoretical masses for all plasmids studied (Table 1, under DNA optimized MP measurement). Next to the shift in mass values, the FWHM of the pUC18, pBR322, and ΦX174 mass distributions are reduced by over two-fold following FA incubation (Fig. 4A), ranging ultimately between ∼160 to 220 kDa. The improved mass accuracy and precision most likely stem from the dramatic drop in the signals’ ellipticity values (Fig. S2). Based on the measurements with the FA-transformed plasmids, we constructed a new MP standard calibration curve. This calibration curve is now linear and aligns closely with the ‘ideal’ curve where the measured mass (based on a protein standard) equals the theoretical mass (Fig. 4B). With the intact plasmids studied here our analyses were limited to a maximum of ∼5 MDa, however, when incorporating data obtained for the multimers of pBR322 and p8064, formed at elevated concentrations of FA (Fig. 3B), we can extend this calibration curve up to ∼15 MDa (Fig. 4C). Although the multimer peaks in this mass range are less populated and broader compared to the detected single plasmid particles, their peak width remains remarkably narrow, especially for pBR322 (Fig. 3B). This indicates that the size and polarizability of the multimers are within limits for ‘conventional’ MP detection based on a protein standard. With the multimers in line with the protein contrast-to-mass conversion, we can potentially calibrate the mass photometer for ultra-high molecular weight particles up to ∼15 MDa (Fig. 4C).

**Figure 4:**
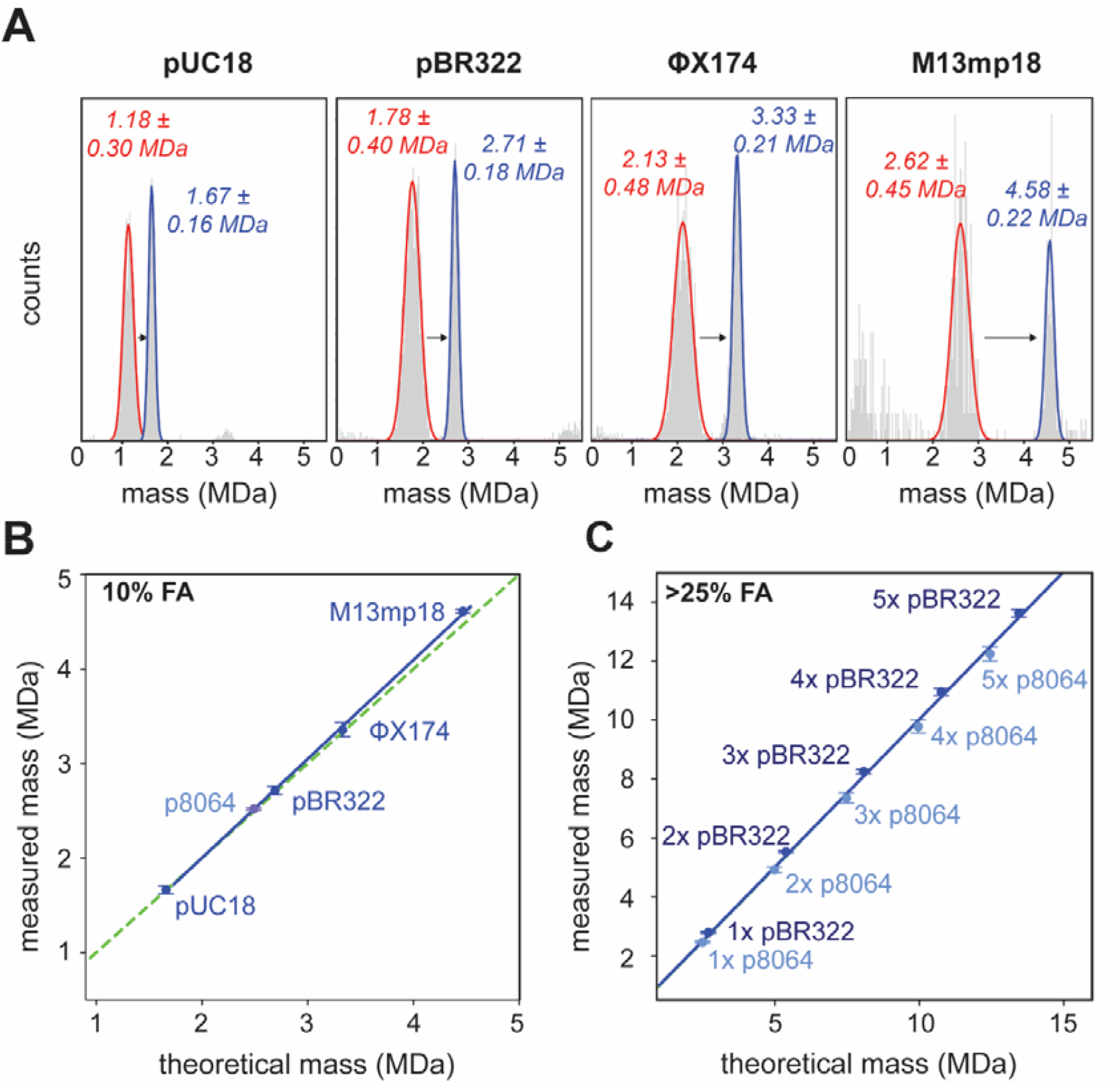
MP of several dsDNA plasmids before and after transformation into ssDNA like particles. **A)** MP measurements of dsDNA plasmids prior to and after incubation with 10% FA for 30 s. An alike characteristic shift in contrast and retrieved mass value is observed for every pDNA construct. Only after incubation with FA did the measured masses fit the expected masses. Depicted are the average mass ± FWHM of the Gaussian fit, measurement without FA indicated in red and with FA indicated in blue. **B)** Following transformation induced by 10% FA the measured masses match the expected masses for all pDNAs studied (the dashed green line represents the perfect match). **C)** By incorporating data from the FA-induced multimers of pBR322 (dark blue) and p8064 (light blue) the calibration curve can be extended further in the ultra-high mass range.

## Discussion

Naked plasmid DNA as a base for a vaccine has picked up interest recently with several newly approved clinical applications (e.g., ZyCoV-D, Collategene) and various ongoing clinical trials.^4,27-29^ Over the years, production methods have been improved in response to the high demand for pDNA for use in research and clinical testing. For quality control of pDNA methods such as capillary electrophoresis and nanopore sequencing have been used. However, such techniques can be time and material-consuming, which may be undesirable at an early stage of research and development. Here we present a simple and quick protocol to measure the mass of pDNA by mass photometry which can be used as an additional characterization method during pDNA development. With the introduction of an efficient sample preparation step, we can still use the conventional MP workflow (as optimized for proteins) and are able to accurately measure plasmids masses between 0.5 and 15 MDa.

To come to this protocol, we needed to first consider the prolonged shape of pDNA. Compared to common protein-based samples, pDNA exhibits an exceptionally elongated shape, which we here show impairs its mass analysis by conventional MP. This issue, however, is not only limited to MP. The often side-by-side compared, single-molecule technique of charge-detection mass spectrometry (CDMS) has also been evaluated for its ability to determine the mass of nucleotide particles such as pDNA and RNA.^30-33^ Generally, CDMS on pDNA generates accurate masses and, in addition, gains insight into the plasmid structure with the particles’ charging profile, which is dependent on the plasmid DNA structure (either linear, relaxed or supercoiled).^31^ However, one drawback of this CDMS approach is that shearing of plasmids during electrospray can affect particle detection, limiting the applicability of CDMS to relatively small DNA plasmids.^30,31^ Moreover, a certain level of expertise is required to limit shearing, optimize pDNA detection and process the data.

With MP, an aqueous in-solution based method, shearing of DNA is not an issue and the data can be simply processed. In turn, the elongated shape of pDNA does reach beyond the diffraction limit of the mass photometer, which can impair the measurement of pDNA (see Fig. 1). Also in an earlier study, even short dsDNA strands (∼400 or 600 bps) measured by MP, displayed a broadening of the contrast histograms due to their non-spherical, anisotropic and lengthy structure.^20^ Here, we alleviate this issue by introducing the FA-induced transformation of the dsDNA to highly compacted ssDNA-like particles (with a concurrent decrease in particle size) as a sample preparation step. By doing so even high molecular weight particles (> 6MDa) appear to remain in a linear contrast-to-mass scattering regime. The compacted ssDNA-like particles land perfectly in MP, even on non-modified glass slides, facilitating the extraction of valid contrast values and accurate masses (Fig. 3 and 4).

With MP, we observe that upon incubation with FA, dsDNA plasmids start to obtain ssDNA-like properties, namely binding to non-modified glass surfaces, protein-like polarizability of light and an overall compacted shape, well below that of the MP diffraction limit. Regrettably, elucidating the actual DNA structure remains difficult. Studies employing AFM demonstrated that an ssDNA fragment with a similar number of bases as a dsDNA construct bears a much more dense and intertwined topology.^34-36^ This can be corroborated by a greatly reduced persistence length of ssDNA and the ability to hybridize with itself.^37-39^ With HPLC, we observed that FA-treated dsDNA plasmids are longer retained on C18 material, similar to ssDNA (Fig. 2). In combination with the MP observations that FA-treated plasmids start to adopt ssDNA-like properties (Fig. 3 and 4), we hypothesize that incubation with FA creates a ssDNA-like state of the plasmids.

Using this protocol, applying for 30 s a 10% concentration of FA to a low concentration of pDNA (∼12.5 ng/μL) seems to instantly convert most of the dsDNA plasmid to particles with a ssDNA-like state. Eventually, these compacted plasmid strands, can also form oligomers (Fig. 3B). It should be noted that FA has also been used to hydrolyze DNA, as sample preparation step for the analysis of the base composition. However, this generally requires much higher concentrations and/or incubation at high temperatures. In our measurements we do not observe any degradation of DNA upon FA incubation.

We suggest that by using this protocol, MP may represent a valuable contribution to the field of vaccine development and cell and gene therapy, enabling the accurate and fast mass analysis of pDNA. By using a single plasmid (i.e., pBR322) and FA we can create a standard curve reaching up to the 10-15 MDa range. With FA, an ssDNA-like state is created out of dsDNA enforcing a globular and more isotropic topology similar to protein. Thus, with only a small adjustment to the conventional MP procedure, we can measure large-sized pDNA and resolve mass peaks better compared to traditional agarose gel electrophoresis systems.

## Methods and materials

### Plasmids

Plasmids of a range of different sizes: pUC18 (2686 bps, 1.74 MDa MW according to manufacturer), pBR322 (4361 bps, 2.83 MDa MW according to manufacturer) and ΦX174 (5386 bps, 3.5 MDa MW according to manufacturer) were purchased at Thermo Fisher Scientific (Vilnius, Lithuania). The M13mp18 plasmid (7249 bps) was purchased from New England Biolabs (Ipswich, MA, USA). The p8064 ssDNA plasmid (8064 nt) was bought at Tilibit (München, Germany).

### Preparation of (coated) coverslips for mass photometry

Glass coverslips (Paul Marienfeld GmbH, 24 x 50 mm, 170 ± 5 µm) were prepared as described earlier.^24,40^ Briefly, for APTES coating, slides were prepared by overnight incubation in 100 mM Sulphuric acid (Merck), after which they were washed by serial rinsing with 1x methanol (Biosolve Chimie SARL, HPLC grade), 1x ethanol (Supelco EMSURE), 1x methanol and 1x ethanol to contain them in fresh ethanol finally. Coating was done in 5% APTES (Sigma) in ethanol for 1 hour at room temperature. After coating, coverslips were washed with ethanol and incubated in 6% acetic acid for 30 minutes. Final cleaning was done by rinsing with methanol, sonication in methanol for 5 minutes and a final wash with 1x methanol, 1x isopropanol (Supelco EMSURE) and drying with N2. Non-coated glass coverslips were prepared by serial rinsing with Milli-Q water and HPLC-grade isopropanol and subsequent drying with N2. Once dry, CultureWell gaskets (Grace Biolabs) were placed on the coverslips as container well for MP measurements.

### Mass photometry measurements

MP measurements were performed on a Samux^MP^ mass photometer (Refeyn Ltd.), essentially as described earlier.^24,40^ After mounting the coverslip in the mass photometer, 12 µL of PBS buffer was applied following focusing of the microscope. Each measurement was initiated by the addition of 3 µL of a sample, mixing it with PBS in the well. Landing events were recorded for 60 seconds with 100 frames per second. Masses were retrieved by converting contrast values based on a calibration mixture consisting of thyroglobulin (Sigma T9145) containing monomers (335 kDa), dimers (670 kDa) and tetramers (1340 kDa) using DiscoverMP software (Refeyn Ltd.). DNA plasmids were diluted to 12.5 - 50 ng/µL either in PBS with FA followed by 30 seconds of incubation with subsequent MP measurement or in PBS with direct MP measurement.

### Ellipticity analysis

After MP recording, the landing events were identified with the DiscoverMP software (Refeyn Ltd.), and individual frames were exported. Going through the frames and bypassing the conventional DiscoverMP filtering step, the contours of all the ratiometric signals were determined. An ellipse was fitted for each landing event using in-house Python scripts employing the OpenCV image processing package (Fig. S3). Ellipticity was calculated by dividing the width by the height of the fitted ellipse.

### RP-HPLC analysis

About 2.5 µg of DNA in PBS, either with or without FA (total volume 10 μL), was injected onto an Agilent 1200 Series HPLC System (Agilent Technologies) and passed over an Aeris Widepore XB-C18 3.6 µm 250 x 2.1 mm column (Phenomenex) with a constant flow rate of 0.25 mL/min. The column was pre-equilibrated in buffer A: 100 mM ammonium acetate (Sigma) pH 6.5 and maintained at room temperature. We applied a 40-minute gradient mixing 60% acetonitrile (buffer B) with buffer A, going from 0 to 40% buffer B. The elution of DNA was monitored at 254 nm UV absorption with fractions taken every 15 seconds. Following the gradient, a 100% buffer B wash was applied for 10 minutes. For the RP-HPLC analysis done prior to MP measurements, we performed a run with 7.5 µg DNA incubated with 25% FA for 15 minutes at room temperature. Following the run, fractions were immediately analyzed by MP, which was set up beforehand with APTES-coated slides.

### Gel electrophoresis

Following incubation of 3.75 μg of pUC18 DNA in 25% FA (room temperature, 10 minutes), the mixture was injected onto the RP-HPLC system and separated for dsDNA/ssDNA as described above except for using a 0.35 mL/min flowrate. Fractions of 50 μL were collected out of which 17 μL was mixed with 3 μL gel loading dye (Tilibit) for loading on an agarose gel. A 1% agarose (Sigma) gel was prepared with GelRed (Biotium) staining. Besides the loaded samples, a supercoiled DNA ladder (New England Biolabs) was added. The gel was run for 90 minutes at 100 V and imaged on an Invitrogen iBright 750 Imaging System (Thermo Fisher Scientific).

## Acknowledgements

This research received support for the Netherlands Organization for Scientific Research (NWO) through the Spinoza Award SPI.2017.028 to AJRH.

## Author contribution

EE: conceptualization, investigation, writing original draft, -review and editing, formal analysis and visualization

ED: conceptualization, investigation, writing -review and editing

MT: resources, writing -review and editing

AR: review and editing

MN: review and editing

AJRH: conceptualization, investigation, writing original draft, -review and editing, resources, funding acquisition

