## supplementary data for "Breaking barriers in the sensitive and accurate mass determination of large DNA plasmids by mass photometry"

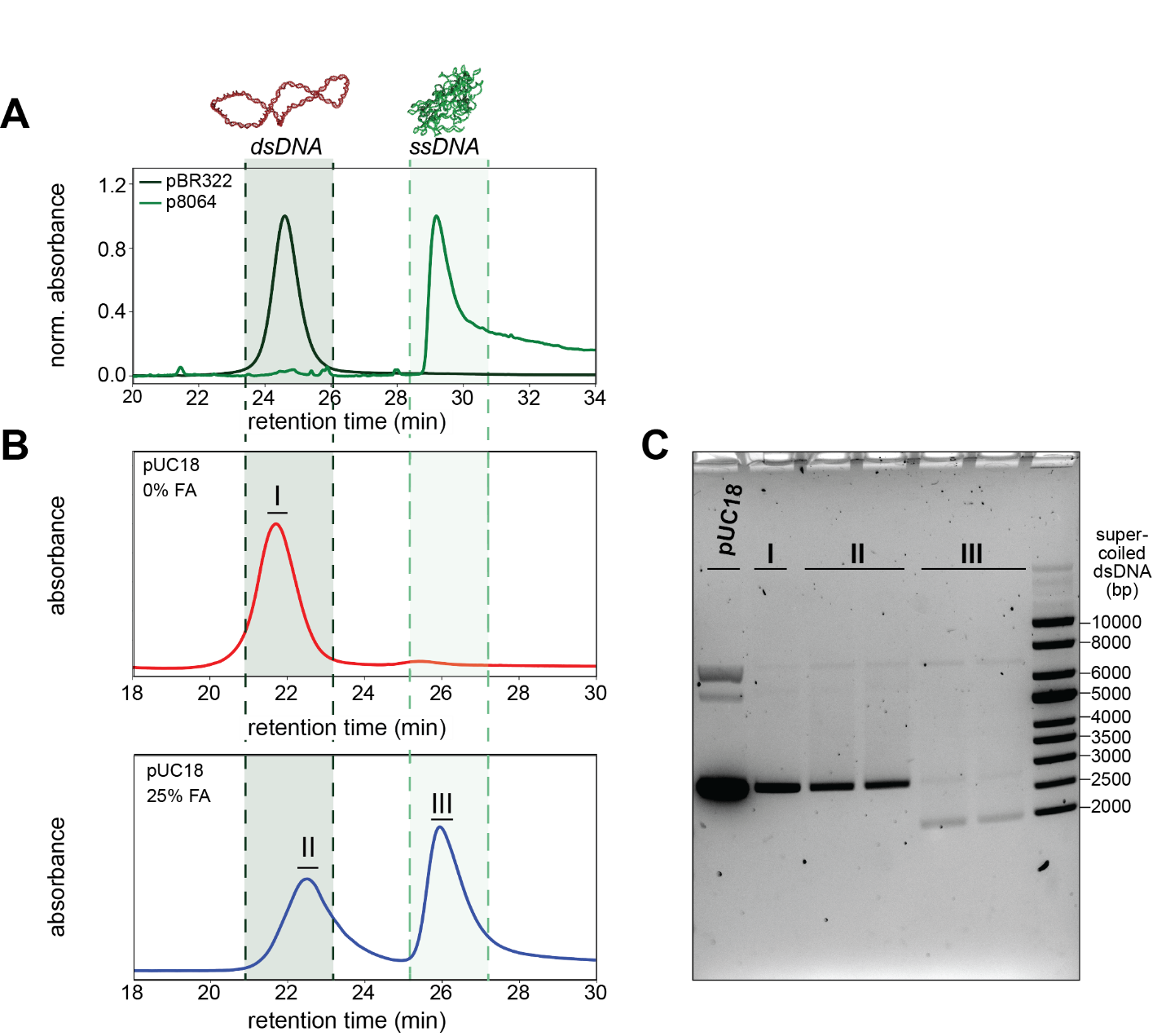

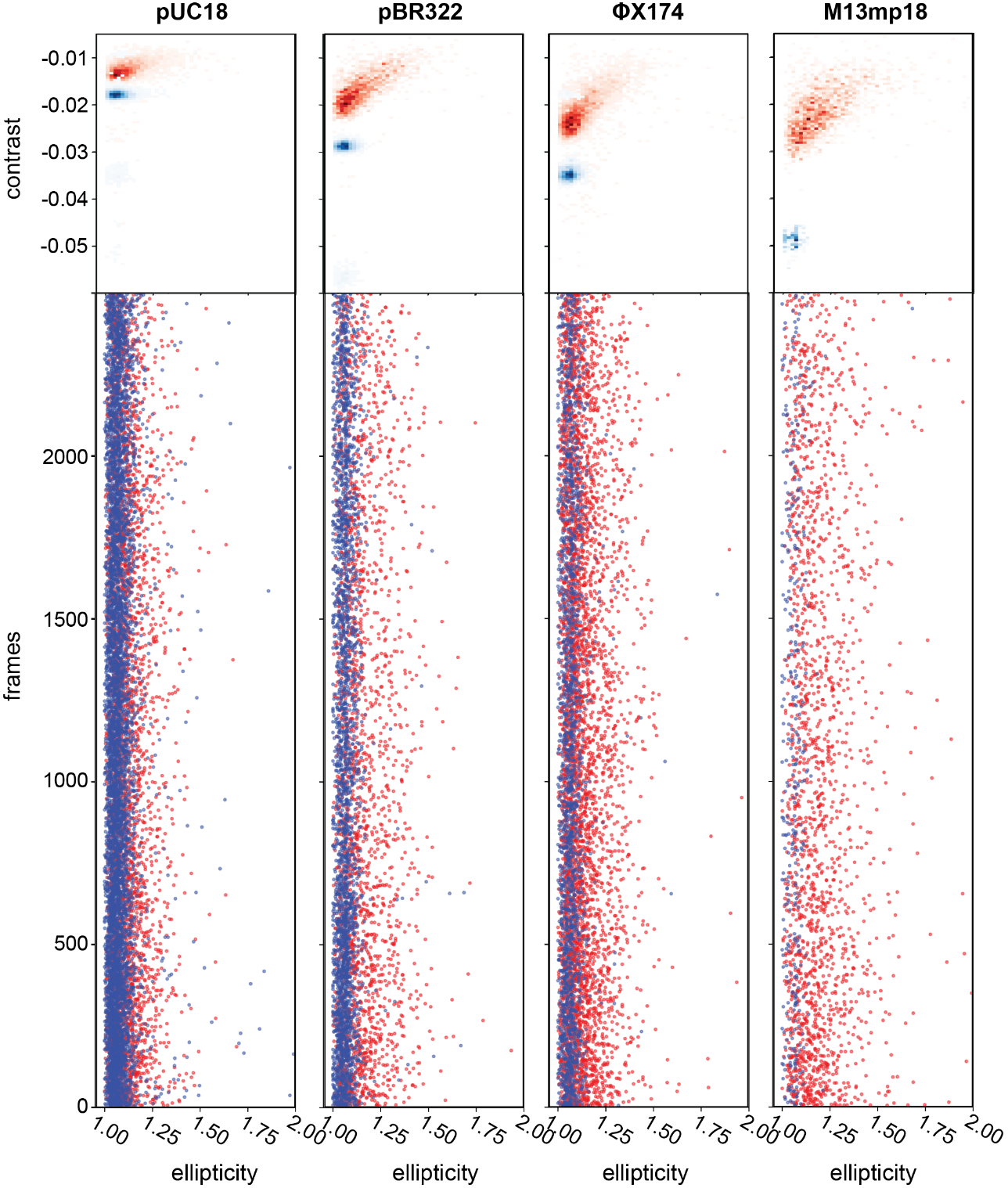

**Figure S1:** **Reversed phase high-pressure liquid chromatography (HPLC) of dsDNA pBR322, pUC18, and ssDNA p8064. A)** Overlayed HPLC chromatograms of similar sized dsDNA pBR322 and ssDNA p8064 (respectively 2.69 MDa and 2.49 MDa), reveal that these separate very well. Using hydrophobic C18 column as stationary phase, pBR322 elutes at ~25 minutes and p8064 at ~29 minutes. **B)** Before (top) and after (bottom) incubation of pUC18 with formic acid (FA), the dsDNA (I and II) and ssDNA-like particles (III) are clearly separated by HPLC. **C)** Fractions were taken for each peak in panel B and analyzed by gel electrophoresis. In fractions I and II, the DNA behaves as dsDNA, in fraction III it behaves like ssDNA running below the dsDNA band.

**Figure S2:** **Characterization of non-ideal landing events for each of the studied pDNA constructs.** Plotted in a 2D histogram are the contrast and ellipticity values of the landing events for the dsDNA plasmids on an APTES-coated glass slide. These landing events were monitored using the native particles (red) or after incubation in 10% formic acid (FA) for 30 s (blue). When not incubated with FA the pDNA particles yield primarily non-circular landing events that impair correct mass determination. Plotting the ellipticity per measurement frame shows that the non-ideal landing events persist throughout the MP recording. Following short incubation with 10% FA the particles’ landing events are mostly circular and show less spread in contrast values, enabling the correct mass analysis by MP.

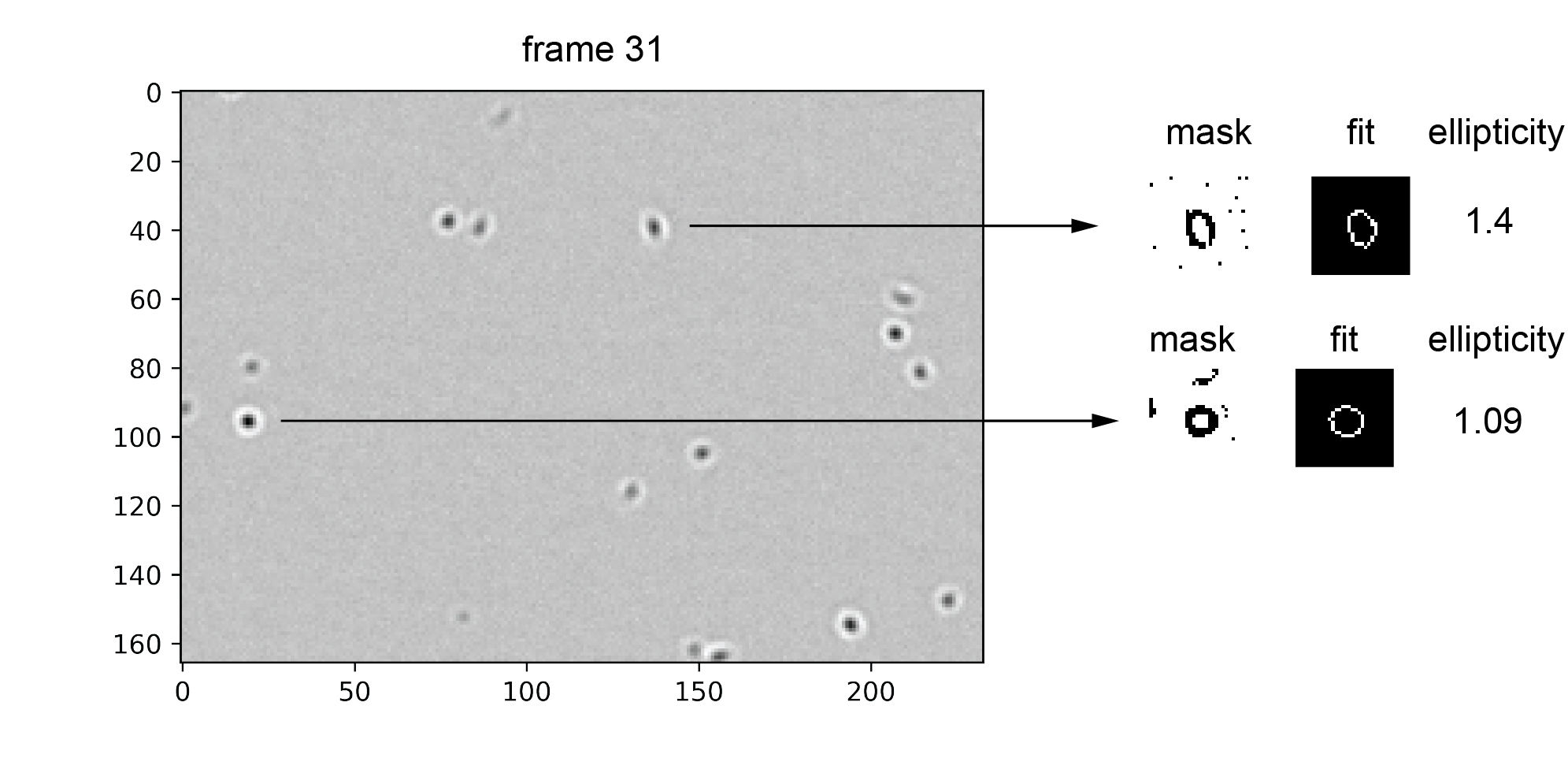

**Figure S3:** **Illustrative example of the extraction of ellipticity values.** Shown is a frame of a pBR322 recording from which two landing events are evaluated for their ellipticity. Following identification of the landing event in the frame, an individual mask is extracted isolating the typical interferometric Airy ring. After fitting, the ellipticity is calculated by dividing the width of the fitted ellipse by its height.
